# Evaluating methods for estimating the proportion of adaptive amino acid substitutions

**DOI:** 10.1101/2022.08.15.504017

**Authors:** Samer I. Al-Saffar, Matthew W. Hahn

## Abstract

A long-standing debate in molecular evolution concerns the role of adaptation in shaping divergence between species. A number of approaches have been developed to estimate the proportion of amino acid substitutions between species (*α*) that are driven by adaptive natural selection. These methods vary in the type of data they use and in the modeling strategies they employ in their inference. In this study, we evaluate the accuracy of nine different methods for estimating *α*, using data simulated in the presence of linked selection. We find that methods that model the distribution of fitness effect (DFE) of both deleterious (as a gamma distribution) and beneficial mutations (as a gamma or exponential distribution) are the most accurate. We applied these methods to whole-genome data, finding that the most accurate methods gave average values of *α*=0.25 in *Arabidopsis thaliana*, 0.5 in *Drosophila melanogaster*, and 0.1 in *Homo sapiens*. We also applied these methods to analyze subsets of tissue-specific genes in *A. thaliana* that are believed to be under different selective pressures and on genes found on the X vs. autosomes in *D. melanogaster*. We find estimates of *α* to be higher in the seeds than in other specialized organs, supporting inferences of conflict-driven adaptive evolution in genes expressed in the seed; we also find *α* to be higher on the X chromosome, supporting previous inferences of faster-X evolution. Overall, our results suggest that there are multiple methods that provide accurate estimates of *α*, providing a guide for future estimates of adaptive evolution.

## Introduction

The role of adaptive mutations in shaping molecular evolution has long been debated (Kimura 1983; Gillespie 1991). With increasing sequence data available from multiple species, it is now evident that natural selection plays a major role in shaping diversity within and between species (Kern and Hahn 2018; Jensen *et al*. 2019). However, questions remain about exactly how much adaptive molecular evolution is needed to explain the patterns we observe, the types of mutations underlying adaptation, as well as the molecular functions most affected by adaptive evolution.

Answering these questions will require many different types of data and many different analytical approaches. Here we focus on tests designed to detect between-species signatures of positive selection on protein-coding genes. One such test was introduced by McDonald and Kreitman (1991). Their test only requires data for a single gene from at least two closely related species, with several individuals sequenced from one of the two species. These data allow us to contrast within-species polymorphism to between-species fixed differences, for both synonymous and nonsynonymous changes. Under neutrality, the ratio of counts of nonsynonymous to synonymous polymorphisms (P_n_/P_s_) should equal the ratio of counts of nonsynonymous to synonymous fixed differences (D_n_/D_s_). Because beneficial mutations are fixed at a higher rate than neutral mutations, their presence will increase D_n_/D_s_ above that of P_n_/P_s_; a statistical test of independence of these values arranged in a 2×2 matrix allows us to reject the null hypothesis of no positive selection (McDonald and Kreitman 1991).

While the original McDonald-Kreitman test allows one to ask about the presence of selection on single protein-coding genes, the terms in this test can also be rearranged to estimate the proportion of excess nonsynonymous substitutions within a gene. Building on the neutrality index of Rand and Kann (1996), Smith and Eyre-Walker (2002) proposed the following measure of the proportion of amino acid substitutions fixed by positive selection:

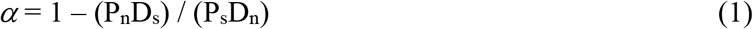

Since the count of polymorphisms and fixed differences can be low or zero at single genes, *α* values are more usefully applied by averaging across many genes, or even whole genomes (Smith and Eyre-Walker 2002; Stoletzki and Eyre-Walker 2011). Using this method, *α* values have been obtained from the whole genome of several species, including *Drosophila melanogaster* (Begun *et al*. 2007), *Arabidopsis thaliana* (Slotte *et al*. 2011), and *Homo sapiens* (Eyre-Walker and Keightley 2009). A major correlate of *α* across species is their effective population size (Galtier 2016), as species with larger effective sizes can fix a larger proportion of slightly advantageous mutations. Studies have also shown a higher proportion of adaptive amino acid substitutions occurring in genes involved in host-pathogen interactions (e.g. Obbard *et al*. 2009; Slotte *et al*. 2011; Enard *et al*. 2016) as well as genes involved in sex-related functions (e.g. Gossman *et al*. 2014).

More sensitive methods for estimating *α*, as well as related evolutionary parameters, are possible when coupling the full site (or allele) frequency spectrum (SFS) to divergence data, rather than just the overall counts of polymorphisms (reviewed in Hahn 2018, chapter 7). Such inferences are possible because both positive and negative selection affect the frequency spectrum in predictable ways (e.g. Sawyer and Hartl 1992). In addition, calculating *α* using just P_n_ and P_s_ will be conservative due to the presence of segregating weakly deleterious variants (Templeton 1996; Fay *et al*. 2002; Charlesworth and Eyre-Walker 2008). Several methods have therefore been developed to estimate *α* using the full site frequency spectrum, with the caveat that non-equilibrium demographic histories and nearby linked selection will perturb the frequency spectrum away from theoretical expectations. Most methods deal with these non-equilibrium scenarios by first fitting a model to the synonymous frequency spectrum—which is assumed to account for the overall effect of linked selection and demography—and then fitting a model to nonsynonymous frequency spectrum taking into account this inferred history in order to estimate selection parameters.

Many different approaches have now been developed to estimate *α* (see Methods), though the results from these methods often differ. One reason for these differences is that the methods are based on models with different underlying assumptions, use different data (e.g. total counts of polymorphisms vs. the allele frequency spectrum), and are implemented using very different computational and numerical approaches. Therefore, here we compare estimates obtained from nine different methods for inferring *α* from whole-genome data. We applied these methods to data simulated across a range of population sizes and selection coefficients, and in the presence of background selection, in order to quantify their accuracy. We also applied these methods to three population genomic datasets obtained from *Drosophila melanogaster, Arabidopsis thaliana*, and *Homo sapiens*. We show how inferences about *α* can vary across methods, highlighting these differences by analyzing subsets of tissue-specific genes in *A. thaliana* that are believed to be under different selective pressures and in a comparison of the X chromosome and autosomes in *D. melanogaster*. Our comparison provides a guide for choosing the appropriate method when estimating adaptive amino acid substitutions from population genomic data.

## Materials and Methods

### Simulated data

To test the accuracy of methods for estimating the proportion of adaptive substitutions, we used a previously published simulated dataset (Booker 2020). Briefly, the simulation package SLiM (Haller and Messer 2019) was used to simulate a Wright-Fisher population of *N* = 10,000 diploid individuals. Each simulated chromosome consisted of seven genes, where each gene consists of five 300 bp exons separated by 100 bp introns; the genes themselves are separated by 8,100 bp of non-coding sequence. In exons, two-thirds of sites were modeled as nonsynonymous and the other third as synonymous.

The rate of mutation (*μ*) and recombination (*r*) were set to 2.5×10^−7^ in the simulation, such that 4*Nr* = 4*Nμ* = 0.01, where *N* is the population size. Advantageous mutations were introduced with two parameters: one specifying the proportion of newly arising advantageous mutations, *p*_*a*_, and another specifying the fitness effects of these mutations, γ_a_ (= 2*Ns*_*a*_, where *s*_*a*_ is the selection coefficient of advantageous mutations). In addition to advantageous mutations, all simulations included constant background selection in coding regions, with the proportion of nonsynonymous deleterious mutations equal to 1- *p*_*a*_. The distribution of fitness effects of deleterious mutations was modeled as a gamma distribution with a mean of γ_d_ = 2*Ns*_*d*_ = −2,000 (where *s*_*d*_ is the selection coefficient of deleterious mutations) and a shape parameter of β = 0.3 (based on estimates from Loewe and Charlesworth 2006). Fifteen different parameter combinations were used to simulate data that vary in the frequency and strength of beneficial mutations (*p*_*a*_ = 0.01, 0.001, and 0.0001; and γ_a_ = 10, 50, 100, 500, and 1000). For each parameter combination, 2000 independent datasets were produced. We merged these replicates for each set of parameters, resulting in a total of 21 Mb of protein-coding sequence for each. Because ancestral states are known, both the unfolded and folded frequency spectrum can be calculated, as can the number of fixed differences.

### Empirical data

We applied methods for estimating *α* to three available whole-genome datasets: one for *Drosophila melanogaster* and one for *Arabidopsis thaliana* (both from Moutinho *et al*. 2019), and one for human (Uricchio *et al*. 2019). Briefly, the *D. melanogaster* data consist of nonsynonymous and synonymous polymorphisms from 110 genomes sampled from 22 sub-Saharan populations. In total, 10,318 genes across all five major chromosome arms were used, with divergence data obtained by comparison to *D. simulans*. For *A. thaliana*, polymorphism data was originally obtained from 18,669 genes within 105 genomes of the Spanish population from the 1001 Genomes Project (Alonso-Blanco *et al*. 2016). Divergence data was obtained by comparison to *A. lyrata*. Lastly, the human polymorphism data was summarized over 16,947 autosomal genes derived from 661 samples representing all seven African subpopulations available from phase 3 of the Human Genome Project (The 1000 Genomes Project Consortium 2015). Human-specific divergence data was obtained by comparison with chimpanzee and orangutan in order to place substitutions along the branch leading to humans. For all three species comparisons, the ancestral state of all mutations was also polarized using the outgroup(s), so that both the unfolded and folded frequency spectrum could be used.

Additionally, we ran analyses separately on sets of genes enriched in different organs of *A. thaliana*. These subsets of genes are described in Geist *et al*. (2019), who tested predictions about adaptive evolution resulting from conflict between maternal and paternal optima within organs (seed, stem, root, floral buds, leaf rosette). We extracted these gene subsets from the Moutinho *et al*. (2019) *A. thaliana* data described above.

### Estimation of *α*

We estimated *α* using nine different methods. The first three methods are based on equation 1, or close approximations of it. We refer to the standard approach using the full counts of P_n_, P_s_, D_n_, and D_s_ as “MK” because these are the same values used in the McDonald-Kreitman test, though this estimator originates with Smith and Eyre-Walker (2002). To avoid the confounding effects of low-frequency deleterious nonsynonymous polymorphisms, Fay, Wyckoff, and Wu (2001) proposed removing all variants (both nonsynonymous and synonymous) at frequencies <15%; we refer to this approach as “FWW.” As slightly deleterious nonsynonymous variants can hitchhike to frequencies above 15%, Messer and Petrov (2013) proposed a method that estimates the expected value of *α* as a function of derived alleles at any frequency, *x*. An exponential function is fit to the data and, as *x* → 1, the value of the asymptote *α*(*x*) should converge close to the true *α* value. Following the original authors, we refer to this method as “asymptotic MK.”

Multiple additional methods for estimating *α* explicitly model the selective effects of deleterious and advantageous nonsynonymous mutations. These methods all require two steps: First, to account for the joint effects of non-equilibrium population demography, linked selection, and other non-selective processes on the frequency spectrum, the SFS of synonymous polymorphisms is fit to a flexible model. Second, given the underlying “demographic” model estimated in the first step, the distribution of fitness effects (DFE) of nonsynonymous mutations is inferred. While there are multiple ways in which step 1 is carried out, the biggest difference among methods is in the shape of the distribution used in step 2. One of the earliest methods fit a gamma distribution to the polymorphism data to estimate the DFE of deleterious mutations only (Eyre-Walker *et al*. 2006; Keightley and Eyre-Walker 2007; Eyre-Walker and Keightley 2009). Subsequent updates to this approach include the effect of beneficial mutations on the SFS: Schneider *et al*. (2011) use a second gamma distribution to model a fraction of the nonsynonymous mutations as beneficial. We refer to these two approaches as “Gamma-Zero” and “Gamma-Gamma,” respectively. Other methods of this type include the “Gamma-Exponential”: in addition to modeling the DFE of deleterious mutations as a gamma distribution, it models beneficial mutations as an exponential distribution. The “Scaled-Beta” method models both weakly deleterious and weakly beneficial mutations as a beta distribution with a flexible shape parameter, where the shape can range from flat to a U-shaped distribution (Galtier 2016). Several methods use approximations of the DFE that have been derived from Fisher’s geometric model (Fisher 1930). These include the “Displaced-Gamma” model (Martin and Lenormand 2006) and the “FGM-BesselK” model (Lourenço *et al*. 2011).

We applied these methods both to the unfolded SFS and, when applicable, to the folded SFS. (Both the “FWW” and “asymptotic MK” methods cannot be used with a folded SFS, and the “MK” result will be exactly the same under the folded and unfolded). In the small number of cases in which the number of polymorphisms at a given sample frequency is zero (which occurs when the *A. thaliana* data are subset into tissue-specific genes), we added a pseudocount of 1 to avoid undefined terms when carrying out calculations.

### Software packages

We estimated the proportion of adaptive amino acid substitutions (*α*) using the following software packages: 1) Grapes (version 1.1): https://github.com/BioPP/grapes (Galtier 2016). This software implements several models, including “MK,” “FWW,” “Gamma-Gamma,” “Gamma-Zero,” “Gamma-Expo,” “Scaled-Beta,” “Displaced-Gamma,” and “FGM-BesselK.” 2) asymptotic MK: http://www.benhaller.com/messerlab/asymptoticMK.html (Haller and Messer, 2017). This program is available as a web-based service, and it implements the method described by Messer and Petrov (2013).

## Results

### Evaluating methods used to infer the proportion of adaptive substitution from simulated datasets

We assessed the accuracy of nine methods to estimate the proportion of adaptive substitutions across a range of simulated datasets by varying parameters for both proportion and fitness effect of advantageous mutations, resulting in 15 parameter combinations (Booker 2020). For each parameter combination, the number of nonsynonymous fixed differences (D_n_) was used to calculate the true value of *α*:

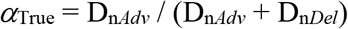

where D_n*Adv*_ is the number of fixed advantageous nonsynonymous differences and D_n*Del*_ is the number of fixed deleterious nonsynonymous differences (Table S1). Each fixed difference in the simulation is labeled as either advantageous or deleterious by SLiM, so these numbers are straightforward to calculate. As expected, there is a strong correlation between the strength of positive selection in each simulation (as measured by the product of *p*_*a*_ and 2*Ns*_*a*_) and the true value of *α* (Spearman’s *ρ*=0.99, *P*<1.0×10^−6^; Table S1). We used the true value of *α* as a baseline for comparison with estimates of *α* obtained from the methods we tested; we call the difference between the two Δ*α*.

We first ran seven methods that use the unfolded (polarized) site frequency spectrum, alongside results from the “MK” method for comparison (Figure 1; Table S2). We found that the “Gamma-Expo,” “Gamma-Gamma,” and “Displaced-Gamma” methods performed very well and showed the highest accuracy, giving average Δ*α* values close to zero. These results suggest that a gamma or exponential distribution is a good fit for inferring the DFE of beneficial mutations for most parameter combinations (Imhoff and Schlotterer 2001; Eyre-Walker *et al*. 2006; Kassen and Bataillon 2006). Note that the data were simulated assuming a gamma distribution of deleterious mutations, but a fixed value (=2*Ns*_*a*_) for all advantageous mutations.

**Figure 1:**
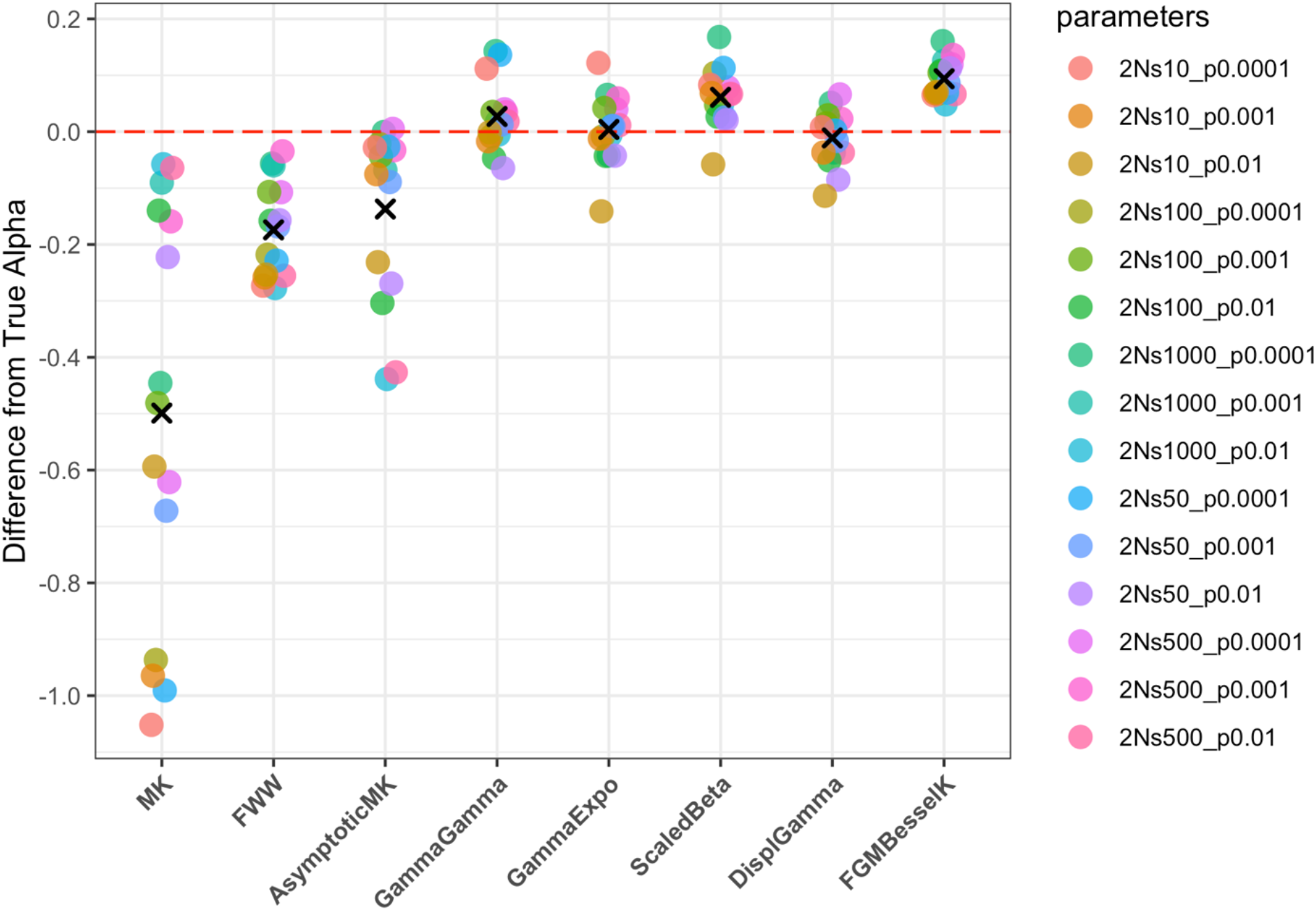
Comparison among *α* values estimated using eight different methods from simulated data. All of the methods (except for “MK”) are using the unfolded site frequency spectrum. The red dashed line represents the true value of *α*, with the y-axis measuring the difference from this value for each estimate. Colored dots indicate different selection parameter combinations used to simulate the data, with the “✕” representing the mean value among parameter combinations. “*2Ns*” is the fitness effect of new advantageous mutations and “*p*” is the proportion of newly arising advantageous mutations.

Results from individual methods largely conformed to expectations. When using the “MK” method (Smith and Eyre-Walker 2002), we found that values of *α* were always underestimates when compared with the true value (Figure 1). In some cases, the difference between the two values is >1, indicating severe inaccuracy. While negative values of the proportion of adaptive substitution are difficult to interpret in a meaningful way, they often result from an excess of nonsynonymous polymorphisms segregating at lower frequencies. The “FWW” method (Fay *et al*. 2001) removes variants segregating at frequencies below 15%, and showed improved results relative to the “MK” method for all tested parameter combinations (Figure 1). However, these results are still all underestimates of the true *α* value. Changing the cutoff for the “FWW” method to exclude variants segregating at frequencies lower than 25% gave similar results (data not shown). We also applied the “asymptotic MK” method (Messer and Petrov 2013), which estimates *α* as a function of derived alleles at any frequency, *x* (where *x* ranges between 0.1 and 0.9). This method gave underestimates of *α* for all but one parameter combination (Figure 1), similar to the previous two methods. Overall, estimates obtained with these three non-parametric methods tend to underestimate *α* values, suggesting that they do not efficiently deal with either the slightly deleterious or slightly advantageous mutations segregating in the sample data, or possibly with the background selection present in the simulations.

Results from methods that estimate *α* by inferring the distribution of fitness effects were much more accurate. The “Gamma-Gamma” method (Eyre-Walker and Keightley 2009; Schneider *et al*. 2011) gave accurate estimates of *α* for all parameter combinations, with the largest difference being <|0.15| from the true *α* value. Similarly, results obtained from the “Gamma-Exponential” method (Galtier 2016) gave estimates of *α* that closely approach the true values for most parameter combinations. By contrast, the “Scaled-Beta” method (Galtier 2016) gave an overestimate for most parameter combinations, though Δ*α* was always less than |0.2|. The last two parametric methods, “Displaced-Gamma” (Martin and Lenormand 2006) and “FGM-BesselK” (Lourenço *et al*. 2011), are based on Fisher’s geometric model of adaptation. For all parameter combinations, results obtained from the “Displaced-Gamma” method were highly accurate (Figure 1), while “FGM-BesselK” overestimated *α* across all simulated datasets.

There was little relationship between any of the results and the values of either *p*_*a*_, 2*Ns*_*a*_, or their product. This outcome is reassuring for two reasons. First, it indicates that methods that give accurate results do so across a range of values of the true *α*, from 1% of all substitutions fixed by positive selection to 95% (Table S1). Second, due to linkage with advantageous nonsynonymous mutations, the SFS among synonymous polymorphisms can be extremely skewed relative to equilibrium expectations. We measured this skew using Tajima’s *D* (Tajima 1989), finding that this statistic taken from each simulated dataset was strongly correlated with the strength of positive selection as measured by the product of *p*_*a*_ and 2*Ns*_*a*_ (Spearman’s *ρ*=-0.96, *P*<1.0×10^−6^; Table S1). The fact that estimates of *α* were accurate across these conditions indicates that methods will be robust to varying “demographic” scenarios (see Discussion).

We also estimated *α* using the folded (unpolarized) SFS. Four of these methods can be applied to unfolded data and were used above: “Gamma-Exponential”, “Scaled-Beta”, “Displaced-Gamma”, and “FGM-BesselK.” The fifth method, “Gamma-Zero” (Eyre-Walker and Keightley 2009), can only be applied to folded data. Note that this last model is also implemented in the software DFE-alpha (Keightley and Eyre-Walker 2007), but we found the version implemented in GRAPES to be more numerically stable. Overall, we obtained results using the folded and unfolded spectra that were quite similar (Figure 2; Table S3). The “Gamma-Exponential” and “Displaced-Gamma” models were once again highly accurate, with “Gamma-Zero” giving equally accurate estimates. Both “Scaled-Beta” and “FGM-BesselK” overestimated *α* using the folded spectrum, often by quite a lot (Figure 2). We again found no consistent association between different parameterizations of the simulations and over- or under-estimation of *α*.

**Figure 2:**
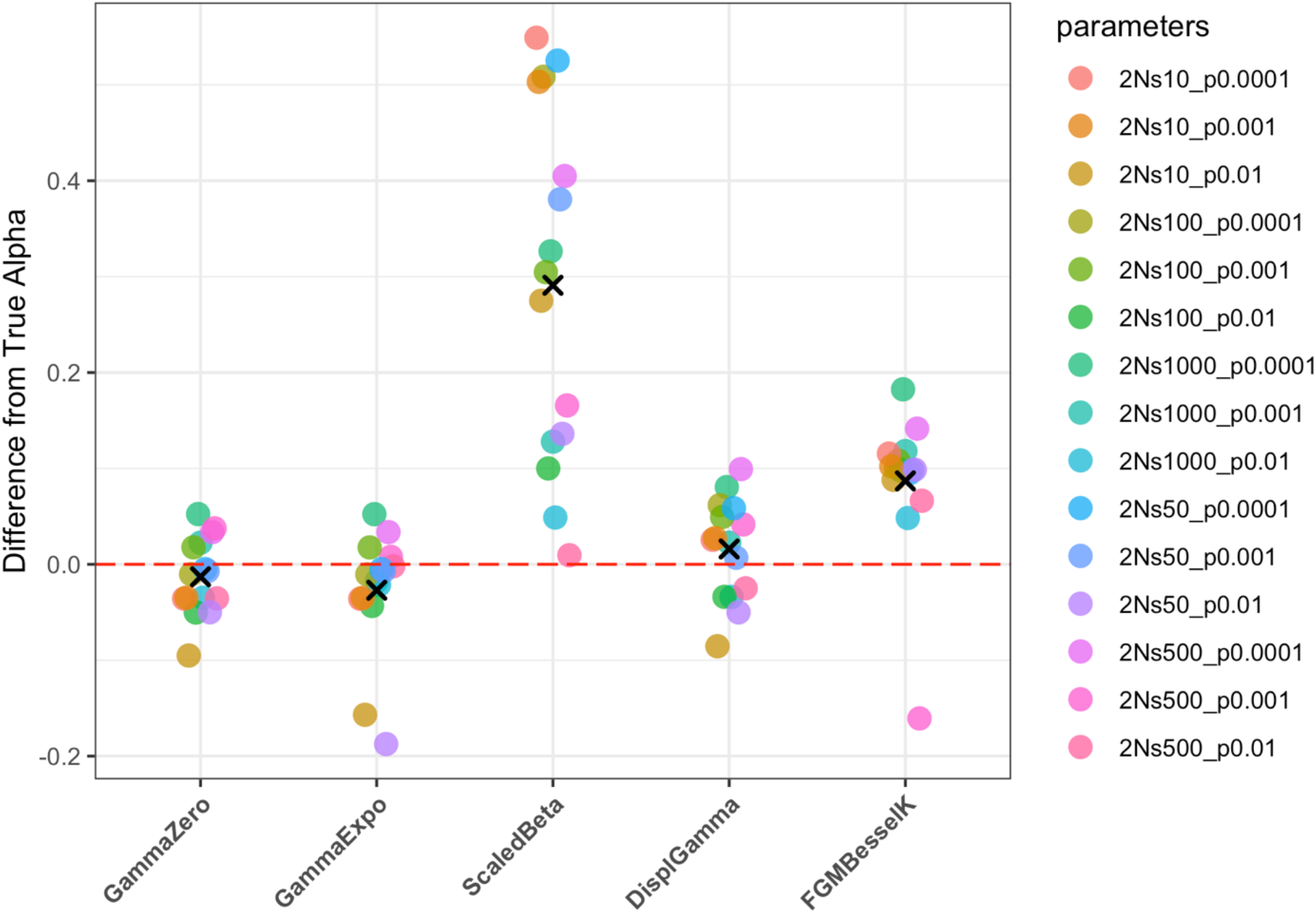
Comparison among *α* values estimated using five different methods from simulated data. All of the methods are using the folded (unpolarized) site frequency spectrum. The red dashed line represents the true value of *α*, with the y-axis measuring the difference from this value for each estimate. Colored dots indicate different selection parameter combinations used to simulate the data, with the “✕” representing the mean value among parameter combinations. “*2Ns*” is the fitness effect of new advantageous mutations and “*p*” is the proportion of newly arising advantageous mutations.

### Adaptive evolution in empirical data

We estimated *α* using all methods applied to whole-genome sequence data obtained from three empirical datasets: *Arabidopsis, Drosophila*, and human (Figure 3). We find trends in the values obtained by different methods on empirical data that are similar to those obtained from simulated data. Methods that do not model the DFE tend to give the lowest values of *α*, often by quite a lot. As expected, the “MK” method gave negative values for all three species, a result that is indicative of the presence of slightly deleterious polymorphisms. The increase in estimates of *α* seen in both the “FWW” and “asymptotic MK” methods demonstrate that fairly straightforward corrections can counteract this problem to some degree.

**Figure 3:**
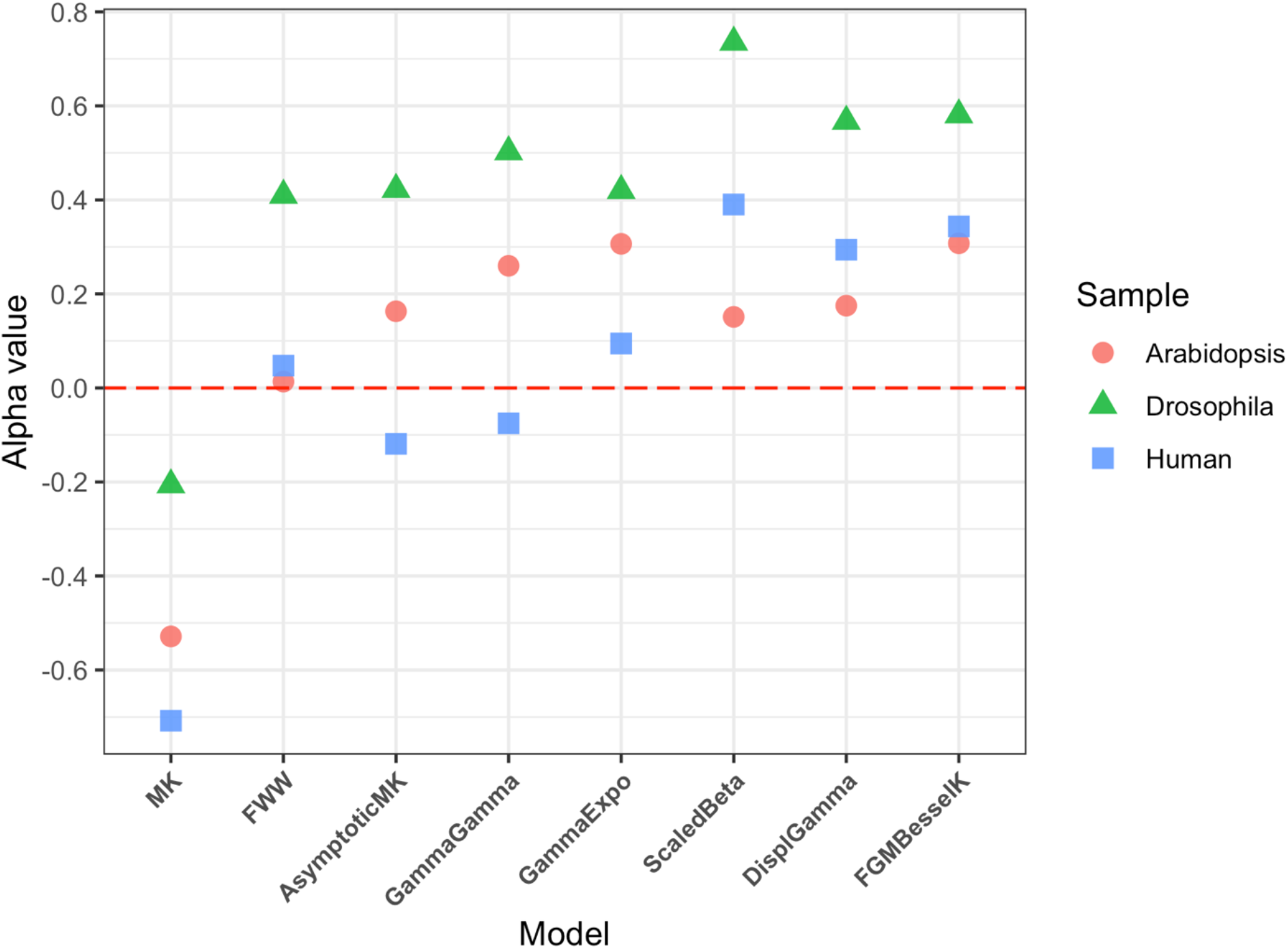
Estimates of *α* values obtained using eight different methods when applied to whole-genome data on the unfolded (polarized) site frequency spectrum from *Arabidopsis thaliana, Drosophila melanogaster*, and *Homo sapiens*. The red dashed line is drawn at a value of 0, while the colors and shapes represent the different species used in this analysis.

Our results from analyses of simulated data suggest that methods that modeling the DFE of mutations can yield more accurate estimates of *α*. Applying these methods to the unfolded biological data gave values of *α* ranging from 0.42-0.74 for *Drosophila*, 0.15-0.31 for *Arabidopsis*, and −0.08-0.39 for human (Figure 3; Table S4). Two of these methods (“Scaled-Beta” and “FGM-BesselK”) tended to overestimate *α* on the simulated data, and were either the two highest estimates (in *Drosophila* and humans) or one of the two highest (in *Arabidopsis*). Among the remaining three methods, the average values of *α* were: *Arabidopsis* = 0.25, *Drosophila* = 0.5, and human = 0.1. We also estimated *α* for the same three datasets using methods that accept the folded SFS as an input (Figure 4; Table S5). In general, we obtained highly similar, but slightly higher, estimates of *α*.

**Figure 4:**
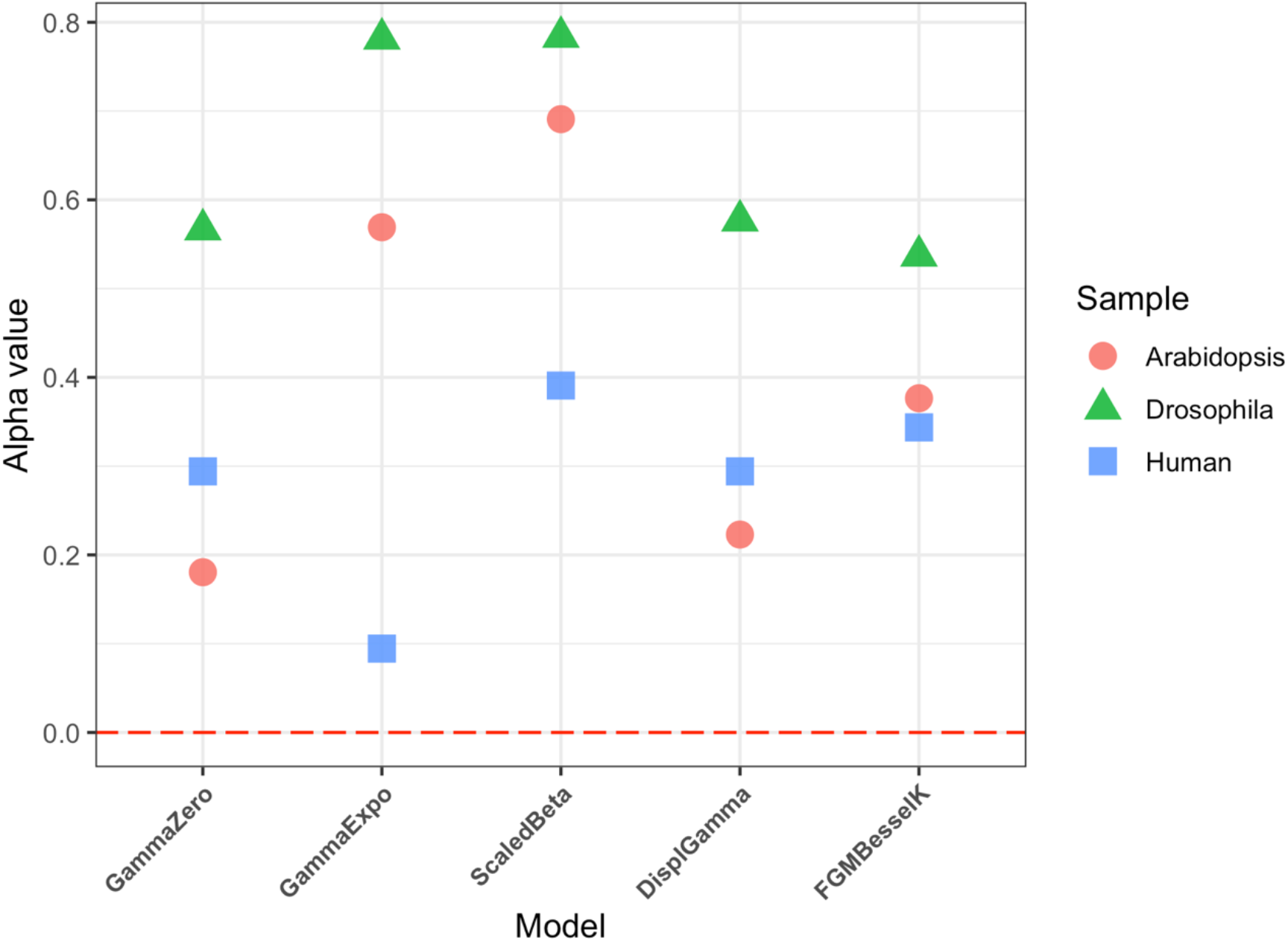
Estimates of *α* values obtained using five different methods when applied to whole-genome data on the folded (unpolarized) site frequency spectrum from *Arabidopsis thaliana, Drosophila melanogaster*, and *Homo sapiens*. The red dashed line is drawn at a value of 0, while the colors and shapes represent the different species used in this analysis.

### Adaptive evolution in different tissues or chromosomes

The proportion of adaptive substitutions is predicted to differ among genes on different chromosomes and among genes involved in different biological processes. For instance, the faster-X effect (Charlesworth, Coyne, and Barton 1987; Meisel and Connallon 2013) posits that genes on the X chromosome will have a greater number of adaptive substitutions. There are a number of possible causes for this pattern, including the exposure of recessive mutations in the heterogametic sex, as well as possible effects of differing recombination and mutation environments. Differences in the function of genes also loosely predicts the amount of adaptive evolution. One consistent pattern is that genes involved in evolutionary conflicts (“arms races”) show a higher proportion of adaptive amino acid substitutions; for instance, this occurs in genes involved in host-pathogen interactions (e.g. Obbard *et al*. 2009; Slotte *et al*. 2011; Enard *et al*. 2016) as well as genes involved in sexual conflict (e.g. Gossman *et al*. 2014).

To test for a faster-X effect in our *Drosophila* data we split the dataset into two parts, one containing genes carried on the X chromosome and the other containing genes carried on the autosomes: 2L, 2R, 3L, and 3R (unfortunately, the human data only include autosomal genes). Calculating *α* using the unfolded SFS, we found higher values for X-linked genes for seven of the eight methods we tested (Table S6). Although the difference between genomic compartments is generally modest, it can sometimes be quite large: using the “MK” method, *α* is estimated to be 0.16 for X-linked genes and −0.3 for the autosomal genes. Our results confirm previous conclusions about the faster-X effect in *Drosophila* (Langley *et al*. 2012), but now using multiple different methods.

To examine *α* in different tissues, we used a large dataset from *A. thaliana*. Geist *et al*. (2019) previously found more adaptive evolution in genes expressed in seeds than in other specialized organs. Because seeds are the arena for conflict between maternal and paternal investment in offspring resources, the genes expressed in this tissue were predicted to show more adaptive evolution. Here we applied all of the methods for estimating *α* to subsets of genes that are preferentially expressed in five different tissues in *Arabidopsis* (Materials and Methods). As found previously, the value of *α* was higher in seeds than in the other four tissues (Figure 5), indicating the presence of more adaptive evolution in genes preferentially expressed in the seed. Within the seed, values of *α* were positive for all methods, with the exception of the “MK” method. Values of *α* varied among tissues, with “floral bud” and “root” also showing generally positive values and “leaf rosette” showing generally negative values. Different methods gave very different estimates of *α* for genes expressed primarily in “stem.” The results for “FGM-BesselK” show positive estimates of *α* across tissues, with an exception in the stem dataset. We believe the negative value in this tissue may be due convergence errors in the optimization function used by the software; we overcame apparently similar such errors in other statistics by running software multiple times, but this negative value of “FGM-BesselK” was found every time.

**Figure 5:**
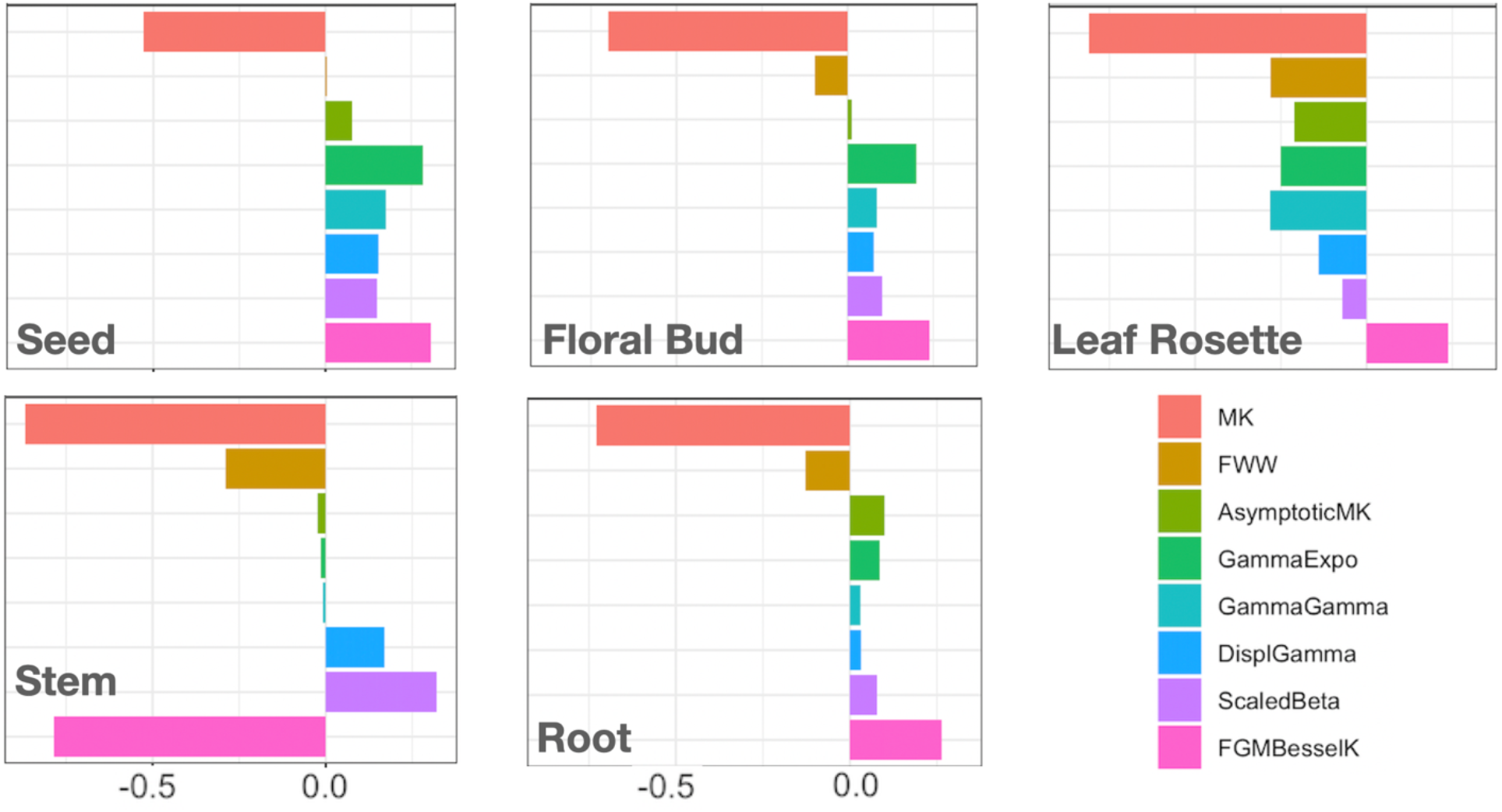
Estimates of *α* values obtained from eight different methods in five different tissues of *Arabidopsis thaliana*. Each panel shows estimates of *α* based on the unfolded site frequency spectrum from genes expressed in that tissue (assignment to tissues was carried out in Geist *et al*. 2018). Within each panel, the values of *α* are centered around zero.

## Discussion

An ongoing debate in evolutionary biology is the relative contribution of selection and drift in shaping genomic diversity among species (Kimura 1983; Gillespie 1991; Kern and Hahn 2018; Jensen *et al*. 2019). One of the widely used approaches for answering this question is to estimate the proportion of fixed nonsynonymous amino acid differences between species, *α*, from different organisms (McDonald and Kreitman 1991; Smith and Eyre-Walker 2002). In this study, we applied nine different methods that estimate *α* to a variety of datasets. Using simulated data, we find that methods that estimate *α* after inferring the DFE of both deleterious and beneficial mutations from the allele frequency spectrum are very accurate, and give more accurate results than methods that do not carry out this step. Interestingly, among these more accurate methods we did not find an advantage to polarizing mutations as either ancestral or derived (i.e. to using the “unfolded” spectrum relative to the “folded” spectrum).

We estimated *α* from whole-genome data for *Arabidopsis thaliana, Drosophila melanogaster*, and *Homo sapiens*. The most accurate methods (as determined by simulation; “Gamma-Gamma,” “Gamma-Expo,” and “Displaced-Gamma”) found that approximately 25% of nonsynonymous changes in the *Arabidopsis* genome have fixed due to positive selection (Figure 3); for *Drosophila* this number is ∼50%, and for human about 10%. These estimates are similar to many estimates from previous studies (e.g. Begun *et al*. 2007; Eyre-Walker and Keightley 2009; Galtier 2016; Uricchio *et al*. 2019; Zhen *et al*. 2021; Geist *et al*. 2019; Moutinho *et al*. 2019). While some older studies in *Arabidopsis* did not find as high a value of *α* (e.g. Slotte *et al*. 2011; Gossmann *et al*. 2014) these were based on data from very early versions of next-generation sequencing technologies. More recent data has consistently shown higher values of *α* (Geist *et al*. 2019; Moutinho *et al*. 2019). Application of these data and methods to test hypotheses about the role of parental conflict in *Arabidopsis* revealed that different methods can give different answers. For instance, in our analysis, the “MK” method gives underestimates of *α* across all tested tissues, and seeds do not stand out as a battleground for parental conflict. However, using all other methods the seeds do have higher estimates of *α* than other tissues.

The simulations used here (originally carried out by Booker 2020) include linked selection, both on advantageous mutations (hitchhiking) and on deleterious mutations (background selection). Linked selection will distort the allele frequency spectrum of nearby polymorphisms (e.g. the values of Tajima’s *D* from synonymous polymorphisms in Table S1) and can both reduce the probability of fixation of weakly beneficial mutations and increase the probability of fixation of weakly deleterious mutations. For these reasons, linked selection may make it more difficult to estimate *α* accurately (Messer and Petrov 2013). In order to avoid such problems, the methods used here that explicitly infer the DFE first fit a “demographic” model to synonymous polymorphisms. Under the assumption that linked selection has homogeneous effects on nearby synonymous and nonsynonymous polymorphisms (Hahn *et al*. 2002)—and that there is no or weak direct selection on synonymous mutations—these methods attempt to infer the distribution of fitness effects on nonsynonymous mutations after taking into account the skew in the frequency spectrum revealed by synonymous polymorphisms. The so-called demographic model that is fit to the synonymous polymorphisms is therefore a complex history of selection and population demography, much as it is for almost all demographic inference from natural populations. The methods considered here that model the DFE appear to do well regardless of the skew in the SFS at linked neutral polymorphisms. Other, more recent, methods have been developed to explicitly account for background selection (e.g. Uricchio *et al*. 2019).

Interestingly, estimates of *α* from these methods are quite similar to those of methods that instead fit more general “demographic” models (e.g. *α*=0.135 for humans using “ABC-MK”; Uricchio *et al*. 2019). In addition, the effect of background selection on linked synonymous sites must be known ahead of time to use these methods. While such methods have the advantage of being able to estimate parameters beyond *α*, data on background selection for non-model species are not readily available, making wider application less feasible.

There is one major assumption made in the simulations we used, and implicitly made by the methods we employ: that the level of selective constraint has been constant over time. This assumption is equivalent to saying that polymorphism data—which by definition reflects relatively recent selective and demographic processes—is an appropriate comparator for divergence data. If, for instance, population sizes were smaller in the past then they are now, a greater proportion of slightly deleterious nonsynonymous mutations will have fixed; as a consequence, we would incorrectly infer these differences as having fixed adaptively (Eyre-Walker 2002; Rousselle *et al*. 2018). Alternatively, selection may have changed because the DFE has shifted between species (e.g. Huber et a. 2017). To account for some potential biases introduced by violations of these assumptions, Tataru *et al*. (2017) proposed a method (“polyDFE”) in which *α* can be estimated from polymorphism data alone without divergence data. However, Booker (2020) found that while *α* was estimated accurately using polyDFE, other selective parameters were misleading and erroneous when advantageous mutations were rare and strongly selected.

With regards to estimates of *α* in the organisms studied here, all three have experienced population expansions over the past tens of thousands of years (e.g. François *et al*. 2008; Tennessen *et al*. 2012; Li and Stephan 2006). Recent expansions can result in an excess of segregating weakly deleterious polymorphism relative to divergence, and therefore do not lead to overestimates of *α*. We therefore consider the estimates obtained here accurate, or even slight underestimates of the true fraction of amin acid substitutions fixed by positive selection. Overall, our results suggest that there are multiple methods that can give accurate estimates of *α*, especially those that explicitly model the DFE of segregating variation. Future estimates of adaptive evolution from non-model species will benefit from using such methods.

## Data availability

No new sequencing data were generated for this study. All data used was previously generated and is publicly available. Simulation datasets were obtained from https://github.com/TBooker/PositiveSelection_uSFS; *Drosophila* and *Arabidopsis* polymorphism and divergence datasets were downloaded from Mountinho *et al*. (2019) supplementary data found at https://academic.oup.com/mbe/article/36/9/2013/5506639#supplementary-data; Human polymorphism and divergence datasets were downloaded from https://github.com/uricchio/mktest/tree/master/data; the *Arabidopsis* list of tissue-specific genes were downloaded from https://github.com/ksgeist/adaptation-in-arabidopsis-seeds/tree/master/Gene_Lists.

## Acknowledgements

We thank Tom Booker, Lawrence Uricchio, David Enard, Ana Moutinho, Julian Dutheil, Katherine Geist, and Nicolas Galtier for sharing data, sharing software, and answering our many questions. Andrew Kern provided helpful feedback on a draft of the manuscript.

## Supplemental Materials

**Table S1:** Values of *α* and Tajima’s *D* obtained from simulating data with 15 different parameter combinations.

| Parameter combination | True $\alpha$ | Tajima's $D^*$ |
| --- | --- | --- |
| 2Ns1000_p0.0001 | 0.4709 | -0.344 |
| 2Ns1000_p0.001 | 0.8316 | -1.684 |
| 2Ns1000_p0.01 | 0.9512 | -1.981 |
| 2Ns500_p0.0001 | 0.3411 | 0.093 |
| 2Ns500_p0.001 | 0.7643 | -1.068 |
| 2Ns500_p0.01 | 0.9331 | -1.893 |
| 2Ns100_p0.0001 | 0.1022 | 0.397 |
| 2Ns100_p0.001 | 0.5004 | 0.058 |
| 2Ns100_p0.01 | 0.8534 | -1.049 |
| 2Ns50_p0.0001 | 0.0533 | 0.411 |
| 2Ns50_p0.001 | 0.3481 | 0.277 |
| 2Ns50_p0.01 | 0.7892 | -0.478 |
| 2Ns10_p0.0001 | 0.0107 | 0.426 |
| 2Ns10_p0.001 | 0.0962 | 0.415 |
| 2Ns10_p0.01 | 0.4999 | 0.304 |

\* These values were calculated only from synonymous polymorphisms

**Table S2:** Mean values for all Δ *α* estimates obtained from full simulation datasets.

| Method | $\Delta\alpha^*$ (Mean) |
| --- | --- |
| MK | -0.499 |
| FWW | -0.174 |
| Asymptotic MK | -0.137 |
| Gamma-Gamma | 0.027 |
| Gamma-Expo | 0.004 |
| Scaled-Beta | 0.061 |
| Displaced-Gamma | -0.011 |
| FGM-BesselK | 0.094 |

\* $\Delta\alpha$ is calculated by subtracting estimated values from true values.

**Table S3:** Mean values for all Δ*α* estimates obtained from folded simulation datasets.

| Method | $\Delta\alpha^*$ (Mean) |
| --- | --- |
| Gamma-Zero | -0.013 |
| Gamma-Expo | -0.027 |
| Scaled-Beta | 0.291 |
| Displaced-Gamma | 0.016 |
| FGM-BesselK | 0.087 |

\* $\Delta\alpha$ is calculated by subtracting estimated values from true values.

**Table S4:** Estimates of *α* obtained from unfolded biological datasets.

| Sample | MK | FWW | Asymptotic-MK | Gamma-Gamma | Gamma-Expo | Scaled-Beta | Displ-Gamma | FGM-BesselK |
| --- | --- | --- | --- | --- | --- | --- | --- | --- |
| Drosophila | -0.21 | 0.41 | 0.42 | 0.50 | 0.42 | 0.74 | 0.57 | 0.58 |
| Arabidopsis | -0.53 | 0.01 | 0.16 | 0.26 | 0.31 | 0.15 | 0.17 | 0.31 |
| Human | -0.71 | 0.05 | -0.12 | -0.08 | 0.09 | 0.39 | 0.29 | 0.34 |

**Table S5:** Estimates of *α* obtained from the folded biological datasets.

| Sample | GammaZero | GammaExpo | DisplGamma | ScaledBeta | FGMBesselK |
| --- | --- | --- | --- | --- | --- |
| Drosophila | 0.57 | 0.78 | 0.58 | 0.78 | 0.54 |
| Arabidopsis | 0.18 | 0.57 | 0.22 | 0.69 | 0.38 |
| Human | 0.29 | 0.09 | 0.29 | 0.39 | 0.34 |

**Table S6:**
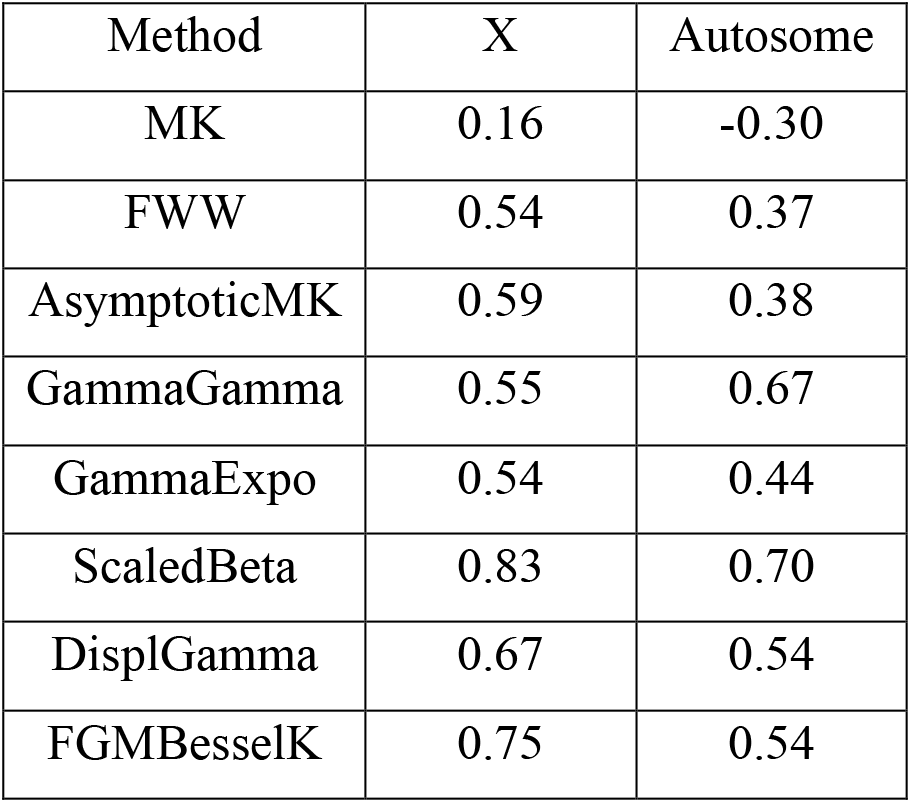
Alpha value estimates obtained from whole-genome data of *D. melanogaster* by dividing the data into two sets: One for genes carried on the X chromosome and the other for genes carried on the autosomes.

## Literature cited

Alonso-Blanco C., J. Andrade, C. Becker, F. Bemm, J. Bergelson, et al., 2016 1,135 Genomes reveal the global pattern of polymorphism in Arabidopsis thaliana. Cell 166: 481–491. https://doi.org/10.1016/j.cell.2016.05.063

Begun D. J., A. K. Holloway, K. Stevens, L. W. Hillier, Y.-P. Poh, et al., 2007 Population genomics: whole-genome analysis of polymorphism and divergence in Drosophila simulans. PLOS Biology 5: e310. https://doi.org/10.1371/journal.pbio.0050310

Booker T. R., 2020 Inferring parameters of the distribution of fitness effects of new mutations when beneficial mutations are strongly advantageous and rare. G3: Genes, Genomes, Genetics 10: 2317–2326. https://doi.org/10.1534/g3.120.401052

Charlesworth B., J. A. Coyne, and N. H. Barton, 1987 The relative rates of evolution of sex chromosomes and autosomes. The American Naturalist 130: 113–146. https://doi.org/10.1086/284701

Charlesworth J., and A. Eyre-Walker, 2008 The McDonald–Kreitman test and slightly deleterious mutations. Molecular Biology and Evolution 25: 1007–1015. https://doi.org/10.1093/molbev/msn005

Enard D., L. Cai, C. Gwennap, and D. A. Petrov, 2016 Viruses are a dominant driver of protein adaptation in mammals. eLife 5: e12469. https://doi.org/10.7554/eLife.12469

Eyre-Walker A., 2002 Changing effective population size and the McDonald-Kreitman test. Genetics 162: 2017–2024. https://doi.org/10.1093/genetics/162.4.2017

Eyre-Walker A., M. Woolfit, and T. Phelps, 2006 The distribution of fitness effects of new deleterious amino acid mutations in humans. Genetics 173: 891–900. https://doi.org/10.1534/genetics.106.057570

Eyre-Walker A., and P. D. Keightley, 2009 Estimating the rate of adaptive molecular evolution in the presence of slightly deleterious mutations and population size change. Molecular Biology and Evolution 26: 2097–2108. https://doi.org/10.1093/molbev/msp119

Fay J. C., G. J. Wyckoff, and C.-I. Wu, 2001 Positive and negative selection on the human genome. Genetics 158: 1227–1234.

Fay J. C., G. J. Wyckoff, and C.-I. Wu, 2002 Testing the neutral theory of molecular evolution with genomic data from Drosophila. Nature 415: 1024–1026. https://doi.org/10.1038/4151024a

Fisher R. A., 1930 The genetical theory of natural selection. Clarendon Press, Oxford (United Kingdom).

François O., M. G. B. Blum, M. Jakobsson, and N. A. Rosenberg, 2008 Demographic history of European populations of Arabidopsis thaliana. PLOS Genetics 4: e1000075. https://doi.org/10.1371/journal.pgen.1000075

Galtier N., 2016 Adaptive protein evolution in animals and the effective population size hypothesis. PLOS Genetics 12: e1005774. https://doi.org/10.1371/journal.pgen.1005774

Geist K. S., J. E. Strassmann, and D. C. Queller, 2019 Family quarrels in seeds and rapid adaptive evolution in Arabidopsis. Proceedings of the National Academy of Sciences 116: 9463–9468. https://doi.org/10.1073/pnas.1817733116

Gillespie J. H., 1991 The causes of molecular evolution. Oxford University Press, Oxford (United Kingdom).

Gossmann T. I., M. W. Schmid, U. Grossniklaus, and K. J. Schmid, 2014 Selection-driven evolution of sex-biased genes is consistent with sexual selection in Arabidopsis thaliana. Molecular Biology and Evolution 31: 574–583. https://doi.org/10.1093/molbev/mst226

Hahn M. W., M. D. Rausher, and C. W. Cunningham, 2002 Distinguishing between selection and population expansion in an experimental lineage of bacteriophage T7. Genetics 161: 11–20. https://doi.org/10.1093/genetics/161.1.11

Hahn M. W., 2018 Molecular population genetics. Oxford University Press, Oxford (United Kingdom).

Haller B. C., and P. W. Messer, 2017 asymptoticMK: A web-based tool for the asymptotic McDonald–Kreitman test. G3: Genes, Genomes, Genetics 7: 1569–1575. https://doi.org/10.1534/g3.117.039693

Haller B. C., and P. W. Messer, 2019 SLiM 3: Forward genetic simulations beyond the Wright–Fisher model. Molecular Biology and Evolution 36: 632–637. https://doi.org/10.1093/molbev/msy228

Huber C. D., B. Y. Kim, C. D. Marsden, and K. E. Lohmueller, 2017 Determining the factors driving selective effects of new nonsynonymous mutations. Proceedings of the National Academy of Sciences 114: 4465–4470. https://doi.org/10.1073/pnas.1619508114

Imhof M., and C. Schlötterer, 2001 Fitness effects of advantageous mutations in evolving Escherichia coli populations. Proceedings of the National Academy of Sciences 98: 1113–1117. https://doi.org/10.1073/pnas.98.3.1113

Jensen J. D., B. A. Payseur, W. Stephan, C. F. Aquadro, M. Lynch, et al., 2019 The importance of the Neutral Theory in 1968 and 50 years on: A response to Kern and Hahn 2018. Evolution 73: 111–114. https://doi.org/10.1111/evo.13650

Kassen R., and T. Bataillon, 2006 Distribution of fitness effects among beneficial mutations before selection in experimental populations of bacteria. Nature Genetics 38: 484–488. https://doi.org/10.1038/ng1751

Keightley P. D., and A. Eyre-Walker, 2007 Joint inference of the distribution of fitness effects of deleterious mutations and population demography based on nucleotide polymorphism frequencies. Genetics 177: 2251–2261. https://doi.org/10.1534/genetics.107.080663

Kern A. D., and M. W. Hahn, 2018 The Neutral Theory in light of natural selection. Molecular Biology and Evolution 35: 1366–1371. https://doi.org/10.1093/molbev/msy092

Kimura M., 1983 The neutral theory of molecular evolution. Cambridge University Press, Cambridge.

Langley C. H., K. Stevens, C. Cardeno, Y. C. G. Lee, D. R. Schrider, et al., 2012 Genomic variation in natural populations of Drosophila melanogaster. Genetics 192: 533–598. https://doi.org/10.1534/genetics.112.142018

Li H., and W. Stephan, 2006 Inferring the demographic history and rate of adaptive substitution in Drosophila. PLOS Genetics 2: e166. https://doi.org/10.1371/journal.pgen.0020166

Loewe, L., and B. Charlesworth, 2006 Inferring the distribution of mutational effects on fitness in Drosophila. Biology Letters 2: 426–430. https://doi.org/10.1098/rsbl.2006.0481

Lourenço J., N. Galtier, and S. Glémin, 2011 Complexity, pleiotropy, and the fitness effect of mutations. Evolution 65: 1559–1571. https://doi.org/10.1111/j.1558-5646.2011.01237.x

Martin G., and T. Lenormand, 2006 A general multivariate extension of Fisher’s geometrical model and the distribution of mutation fitness effects across species. Evolution 60: 893–907. https://doi.org/10.1111/j.0014-3820.2006.tb01169.x

McDonald J. H., and M. Kreitman, 1991 Adaptive protein evolution at the Adh locus in Drosophila. Nature 351: 652–654. https://doi.org/10.1038/351652a0

Meisel R. P., and T. Connallon, 2013 The faster-X effect: Integrating theory and data. Trends in Genetics 29: 537–544. https://doi.org/10.1016/j.tig.2013.05.009

Messer P. W., and D. A. Petrov, 2013 Frequent adaptation and the McDonald–Kreitman test. Proceedings of the National Academy of Sciences 110: 8615–8620. https://doi.org/10.1073/pnas.1220835110

Moutinho A. F., F. F. Trancoso, and J. Y. Dutheil, 2019 The Impact of protein architecture on adaptive evolution. Molecular Biology and Evolution 36: 2013–2028. https://doi.org/10.1093/molbev/msz134

Obbard D. J., J. J. Welch, K.-W. Kim, and F. M. Jiggins, 2009 Quantifying adaptive evolution in the Drosophila immune system. PLOS Genetics 5: e1000698. https://doi.org/10.1371/journal.pgen.1000698

Rand D. M., and L. M. Kann, 1996 Excess amino acid polymorphism in mitochondrial DNA: Contrasts among genes from Drosophila, mice, and humans. Molecular Biology and Evolution 13: 735–748. https://doi.org/10.1093/oxfordjournals.molbev.a025634

Rousselle M., M. Mollion, B. Nabholz, T. Bataillon, and N. Galtier, 2018 Overestimation of the adaptive substitution rate in fluctuating populations. Biology Letters 14: 20180055. https://doi.org/10.1098/rsbl.2018.0055

Sawyer S. A., and D. L. Hartl, 1992 Population genetics of polymorphism and divergence. Genetics 132: 1161–1176. https://doi.org/10.1093/genetics/132.4.1161

Schneider A., B. Charlesworth, A. Eyre-Walker, and P. D. Keightley, 2011 A method for inferring the rate of occurrence and fitness effects of advantageous mutations. Genetics 189: 1427–1437. https://doi.org/10.1534/genetics.111.131730

Slotte T., T. Bataillon, T. T. Hansen, K. St. Onge, S. I. Wright, et al., 2011 Genomic determinants of protein evolution and polymorphism in Arabidopsis. Genome Biology and Evolution 3: 1210–1219. https://doi.org/10.1093/gbe/evr094

Smith N. G. C., and A. Eyre-Walker, 2002 Adaptive protein evolution in Drosophila. Nature 415: 1022–1024. https://doi.org/10.1038/4151022a

Stoletzki N., and A. Eyre-Walker, 2011 Estimation of the neutrality index. Molecular Biology and Evolution 28: 63–70. https://doi.org/10.1093/molbev/msq249

Tajima F., 1989 Statistical method for testing the neutral mutation hypothesis by DNA polymorphism. Genetics 123: 585–595. https://doi.org/10.1093/genetics/123.3.585

Tataru P., M. Mollion, S. Glémin, and T. Bataillon, 2017 Inference of distribution of fitness effects and proportion of adaptive substitutions from polymorphism data. Genetics 207: 1103–1119. https://doi.org/10.1534/genetics.117.300323

Templeton A. R., 1996 Contingency tests of neutrality using intra/interspecific gene trees: The rejection of neutrality for the evolution of the mitochondrial cytochrome oxidase II gene in the hominoid primates. Genetics 144: 1263–1270. https://doi.org/10.1093/genetics/144.3.1263

Tennessen J. A., A. W. Bigham, T. D. O’Connor, W. Fu, E. E. Kenny, et al., 2012 Evolution and functional impact of rare coding variation from deep sequencing of human exomes. Science 337: 64–69. https://doi.org/10.1126/science.1219240

The 1000 Genomes Project Consortium, 2015 A global reference for human genetic variation. Nature 526: 68–74. https://doi.org/10.1038/nature15393

Uricchio L. H., D. A. Petrov, and D. Enard, 2019 Exploiting selection at linked sites to infer the rate and strength of adaptation. Nature Ecology & Evolution 3: 977–984. https://doi.org/10.1038/s41559-019-0890-6

Zhen Y., C. D. Huber, R. W. Davies, and K. E. Lohmueller, 2021 Greater strength of selection and higher proportion of beneficial amino acid changing mutations in humans compared with mice and Drosophila melanogaster. Genome Research 31: 110–120. https://doi.org/10.1101/gr.256636.119

